# Profiling multi-body interactions of BCL2 with single-molecule co-immunoprecipitation reveals the molecular mechanism of ABT-199 resistance

**DOI:** 10.1101/2024.02.04.578783

**Authors:** Changju Chun, Tae-Young Yoon

## Abstract

A capability to characterize protein-protein interactions (PPIs) in a quantitative manner with an increased speed would form a technical basis for accelerating drug discovery targeting the PPI network. We here used the single-molecule pull-down and co-IP platform to examine PPI between BCL2 and BH3-only proteins in crude extract environments. We focused on how the PPI strengths changed with single-point BCL2 mutations found in relapsed chronic lymphocytic leukemia showing ABT-199 resistance, where we took a mix-and-match approach to examine various pairs of baits and preys while titrating their concentrations. This allowed us to examine total 21 PPI reactions and 420 data points, forming a high-resolution large data set of the dissociation constants (*K*_d_) and the drug inhibitory constants (*K*_i_). Our data suggest that the different BCL2 mutants take different routes to acquire resistance to ABT-199, demonstrating how large-scale, quantitative PPI data sets reveal insights into the evolving dynamics of PPI networks.

## Introduction

Protein-protein interactions (PPIs) form the intricate web through which cellular information is transmitted.^1-3^ Selectively amplifying or diminishing specific PPI subsets can alter these networks’ equilibrium, thereby adjusting the flow of signals across various pathways and enabling cells to navigate ever-changing environments. The fine-tuning of PPI intensities is facilitated by various biochemical mechanisms, such as post-translational modifications, scaffolding proteins, and the condensation of intrinsically disordered regions.^4-6^ A quantitative assessment of the strengths of these protein-protein connections, many involving complex multi-body interactions, would enhance our comprehension of PPI networks and their adaptive evolution.

Currently, surface plasmon resonance (SPR) is the benchmark for quantifying PPIs, offering label-free interaction monitoring.^7^ However, SPR signals can be challenging to decipher, especially with multi-body interactions, because the absence of labels complicates the identification of signal origins.^8^ Additionally, SPR typically demands large sample volumes due to measurements at various concentration levels and the necessity of continuous sample flow, resulting in lengthy assessment times for each PPI pair.^8-9^ Therefore, there is a significant opportunity for advancements in high-throughput techniques capable of characterizing multi-body interactions involving multiple protein and small molecule pairs.

Apoptosis is an evolutionarily conserved form of programmed cell death that is important for removing abnormal cells such as cancer cells while causing minimal damage to surrounding tissues.^10-11^ The BCL2 protein family regulates initiation of the apoptosis, and the delicate balance between the various PPIs among its members determines cell fate.^12^ The primary effectors of apoptosis, BAX and BAK, form pores on mitochondrial outer membranes upon undergoing conformational changes triggered by the binding of BH3-only proteins, such as BIM, BAD, and BID.^13-17^ Another group of BCL2 proteins, which includes BCL2 itself, as well as BCLxL and MCL1, interacts with both these effectors and the BH3-only proteins.^12, 18-19^ In many cancers, PPIs involving anti-apoptotic proteins are found to be upregulated than is found in normal cells,^19-23^ effectively preventing interactions between the effectors and the pro-apoptotic BH3-only proteins and thereby preventing apoptosis initiation.^24-26^

Therapeutic agents, referred to as BH3 mimetics, structurally emulate the alpha-helical BH3 domain and are designed to bind anti-apoptotic proteins. This binding effectively releases and/or prevents any further binding with pro-apoptotic proteins,^27-28^ making BH3 mimetics a salient example of a PPI inhibitor.^29-31^ Indeed, ABT-199, which specifically inhibits the PPIs of BCL2 and not BCLxL or MCL1,^30^ was approved for the treatment of chronic lymphocytic leukemia (CLL) in 2016 (NCT01889186), and acute myeloid leukemia (AML) in 2018 (NCT02203773 and NCT02287233). Despite a remarkable initial response rate in phase I trials for relapsed/refractory CLL or small lymphocytic leukemia (SLL), 27% of patients showed regression within 15 months of ABT-199 treatment,^32-33^ often associated with BCL2 mutations such as G101V.^34-35^ The aspartate residue at position 103, which lines the binding groove of BCL2, was also found to be a hotspot, with various single residue substitutes, including tyrosine (D103Y), glutamine (D103E), and valine (D103V) found in relapsed CLL patients.^36-37^ This illustrates how single point mutations in BCL2 can disrupt PPIs between various BH3-only proteins and effectors to affect the balance of the whole PPI network. It also provides an example that underscores the need for quantitative characterization of PPI networks on a larger scale.

Single-molecule pull-down and co-IP (SMPC) is a platform that can assess PPI strength under crude lysate conditions.^38-39^ In this platform, bait proteins are immobilized on a surface via antibody binding. Then, prey proteins labeled with enhanced green fluorescent proteins (eGFPs) are added and allowed to freely diffuse and interact with bait. The formation of a single PPI complex between bait and prey can then be detected as a stable diffraction-limited fluorescence spot under a total internal reflection (TIR) fluorescence microscope with single-molecule resolution.^38-39^ Counting these single PPI complexes within a fixed area provides a rapid way to quantify PPI strength. SMPC has been used to examine the PPI networks of various isoforms of RAS proteins,^40^ epidermal growth factor receptors,^41^ as well as their heterodimer complexes,^42^ all of which work as oncoproteins (or onco-complexes) driving the progression of different types of cancers. In particular, the PPI strengths determined by SMPC have proven to have highly quantitative correlations with the effectiveness of targeted drugs.^41^

Here, we employed the SMPC platform for a quantitative examination of PPIs between BCL2 and BH3-only proteins. Taking advantage of the robustness and speed of the SMPC platform, we generated a large-scale data set encompassing 420 different PPI reactions, including those between BCL2 containing single-point mutations found in relapsed CLL patients and various splice variants of BH3-only proteins. By titrating prey and drug concentrations, we were able to determine dissociation constants (*K*_d_) of PPI and drug inhibitory constants (*K*_i_) for ABT-199. With the increased throughput and quantitativeness provided by this technique, we were able to identify the changes in the *K*_d_ and *K*_i_ values for each interaction with the various BCL2 mutants. Our results point to the different routes by which BCL2 mutations can lead to ABT-199 resistance.

## Results and Discussion

### Application of SMPC to the BCL2 PPI network

To investigate the PPI network with a focus on the anti-apoptotic BCL2 protein, we employed the SMPC platform. We engineered a fusion of a red fluorescent protein (RFP, specifically mCherry) to the C-terminus of the *BCL2* gene, promoting transient expression of BCL2-mCherry in HEK293T cells (Figure 1A). Given that BCL2 proteins possess a single transmembrane domain (TMD),^43^ we extracted the fluorescently labeled BCL2 using mild cholesterol-mimicking detergents, as our prior work validated this method for preserving surrounding membrane patches.^42^ Anti-RFP antibodies, layered atop polyethylene glycol (PEG) polymers, allowed us to immobilize BCL2 proteins directly from cell extracts onto a surface while minimizing the nonspecific binding of other proteins (Figure 1A and Figure S1). This strategy leverages the absence of mCherry homologs in mammalian proteomes to reduce the risk of nonspecific immunoprecipitation. In separate cultures, we expressed various BH3-only proteins tagged with eGFPs (Figure 1B). Lysis of these cells followed by dilution produced extracts with a consistent concentration (typically 10 nM) of the targeted BH3-only protein. These extracts were then introduced into the reaction chamber to initiate PPIs (Figure 1B).

**Figure 1.**
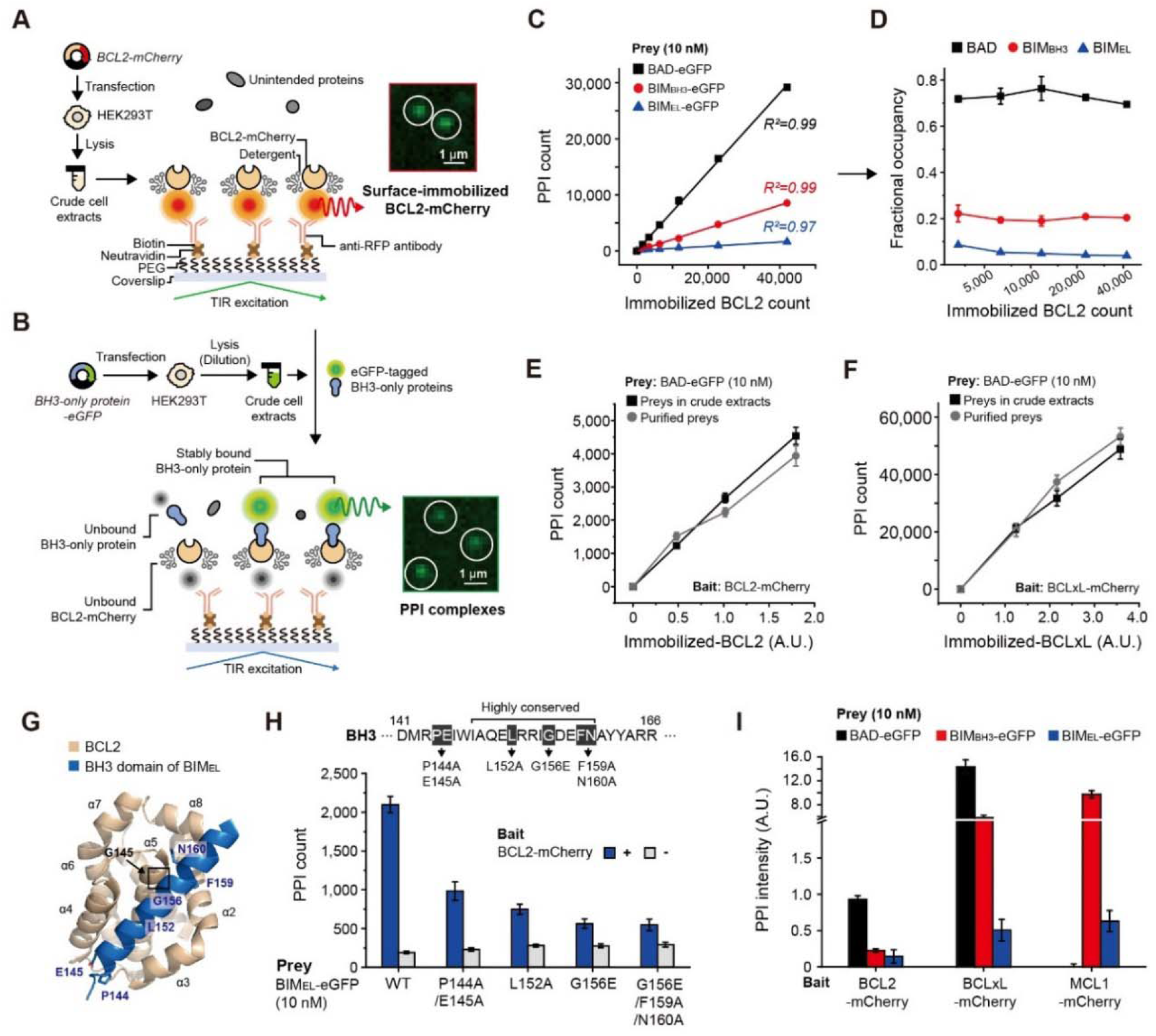
Application of SMPC platform to measure PPIs of BCL2 family members. (A) Schematic for the surface immobilization of mCherry-labeled bait proteins (BCL2-mCherry) overexpressed in HEK293T cells. Bait proteins were selectively immunoprecipitated with a biotinylated anti-RFP antibody. Immobilized bait proteins were observed as defined spots of fluorescent signal (inset). (B) PPIs were initiated by adding eGFP-labeled BH3-only proteins (BAD, BIM_BH3_, and BIM_EL_) overexpressed in HEK293T cells. Prey proteins selectively interacted with immobilized bait proteins. PPI complexes appeared as distinct spots of fluorescent signal (inset). (C) Linear increase in the PPI count with increasing amounts of surface-immobilized BCL2 in the presence of 10 nM of eGFP-labeled BH3-only proteins. (D) Remaining of fractional occupancies with increasing amounts of surface-immobilized BCL2. (E and F) Comparison of the PPIs by the BAD preys in crude extracts and the purified BAD preys. (E) BCL2-BAD PPIs, (F) BCLxL-BAD PPIs. (G) Structure of the BCL2-BIM_BH3_ complex as predicted by AlphaFold2. Highly conserved residues on the BH3 domain of BIM are labeled. G156 on the BH3 domain is located adjacent to G145 on BCL2 (inset). (H) Changes in the PPI count between BCL2 and BH3 domain mutated BIM_EL_. (I) PPIs for various pairs of mCherry-labeled anti-apoptotic proteins (BCL2, BCLxL, and MCL1) and eGFP-labeled BH3-only proteins (BAD, BIM_BH3_, and BIM_EL_).

Single-molecule fluorescence imaging, averaged over hundreds of milliseconds, effectively eliminated the signal from unbound eGFP-tagged proteins due to rapid diffusion, resulting in a diffuse background. In contrast, eGFP-labeled proteins bound to immobilized BCL2 molecules produced sharp, diffraction-limited spots (Figure 1B, inset). We quantified these discrete PPI complexes across approximately 12 fields of view (50×100 µm2) to bolster the statistical reliability of our data. This quantification process, facilitated by an automated focusing system and a robotic stage on our SMPC setup, is completed within a minute (Figure S1).

Upon introducing different pro-apoptotic BH3-only proteins as preys at a concentration of 10 nM, the number of PPIs rose linearly with the increasing number of immobilized BCL2 proteins per field of view (up to 40,000) (Figure 1C). These findings demonstrated that the BCL2 proteins’ fractional occupancy remained essentially unchanged, even with an ∼20-fold increase in BCL2 pull-down numbers (Figure 1D). This constancy implies that the prey proteins in the reaction chambers were not depleted and could sustain equilibrium reactions, even at the highest surface densities tested (approximately 8 baits per μm^2^, as shown in Figure 1D). Consequently, the occupancy values ascertained through SMPC could be directly applied to calculate the dissociation constant (*K*_d_) values for individual binding reactions (see below).

Subsequently, we aimed to discern the impact of endogenous proteins in the extracts on reconstituted target PPI reactions. Employing the protocol described above, we overexpressed BAD proteins in HEK293T cells and diluted the resultant extract to achieve a target BAD protein concentration of 10 nM (Figure S2A). Repeated trials produced diluted extracts with total protein concentrations (*C*_tot_) near 0.1 mg/ml, a dilution exceeding 1,000-fold from native cellular conditions. Adding BAD-eGFP proteins in crude extracts as preys, we found the BCL2 and BCLxL surface baits’ occupancy remained consistent, mirroring results from Figure 1D (Figure 1E, F, black). Remarkably, when comparing PPI reactions utilizing preys in crude extracts with those incorporating purified preys at 10 nM, the binding numbers and occupancy rates for the BCL2 and BCLxL baits were identical (Figure 1E, F, and Figure S3).

These results imply that overexpression, when effectively achieved, renders endogenous proteins negligible in their effect on reconstituted target PPI reactions. We further supplemented the prey protein solution with dense HEK293T cell extracts, investigating alterations in binding counts within more concentrated extract conditions (Figure S2A). Intriguingly, 90% of the binding was retained even with *C*_tot_ in the extract raised to 0.4 mg/ml in extract conditions (Figure S2B, C). We then varied the total protein concentration of the extract for weaker PPI reactions between BCL2 and BIM variants, which are characterized by *K*_d_s in the tens of nM range (see Figure 2 below). Despite the anticipated increased competition from endogenous proteins due to the weaker interaction, the binding count consistently remained at 90% up to *C*_tot_ of 0.4 mg/ml (Figure S2D-F). Accordingly, our proposed method for analyzing PPIs using the SMPC platform has proven to generate reliable PPI counts, even amidst the challenges of a dense extract milieu.

**Figure 2.**
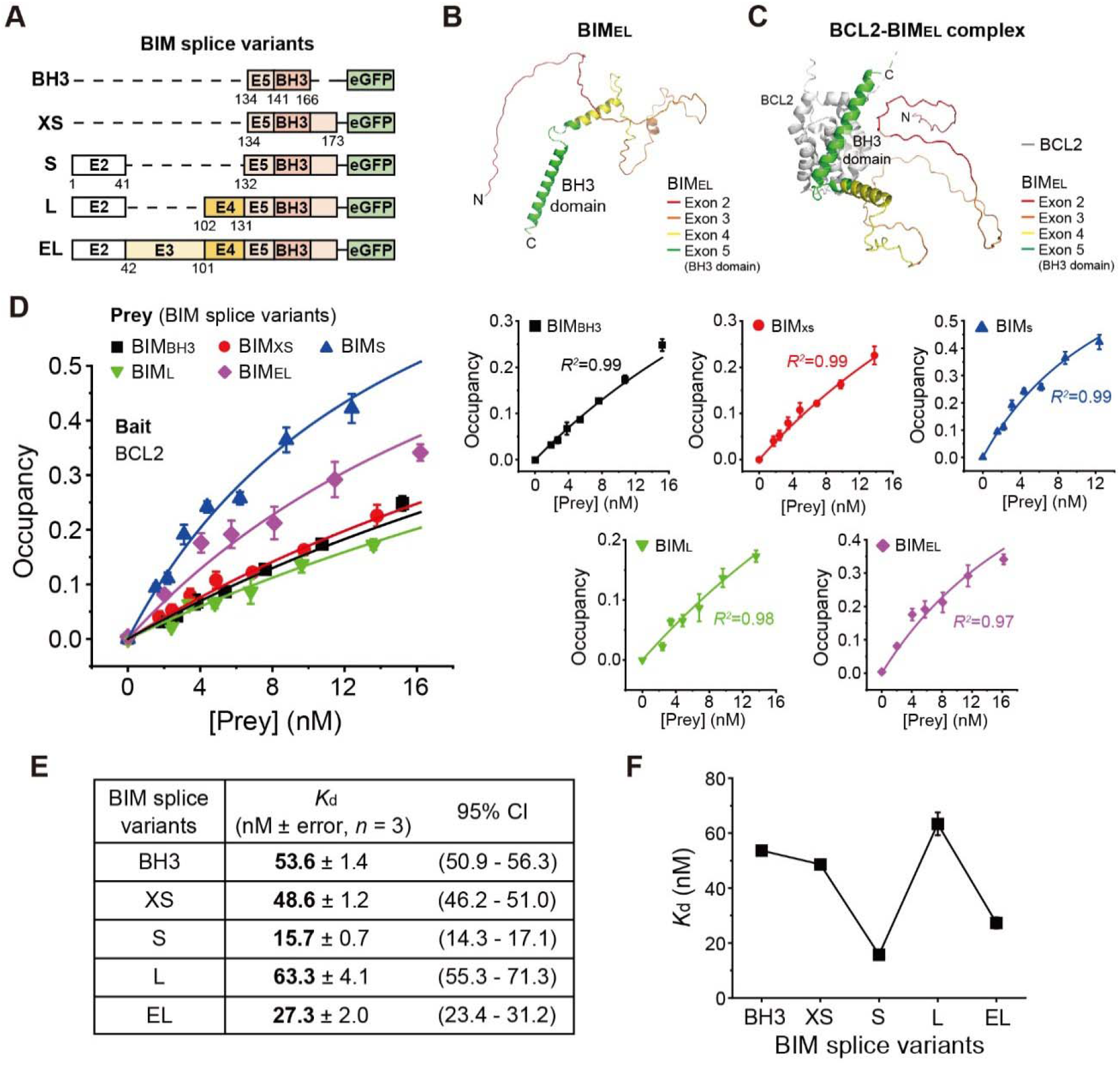
N-terminal exon domains of BIM make distinct contributions to BCL2 PPIs. (**A**) TMD-truncated BIM splice variant (BIM_BH3_, BIM_XS_, BIM_S_, BIM_L_, and BIM_EL_) constructs. E2-E5 refer to each exon domain. (**B**) Predicted structure of BIM_EL_ by AlphaFold2. TMD sequence of BIM was removed for structure prediction. (**C**) Predicted structure of BCL2-BIM_EL_ by AlphaFold2. TMD sequences of BCL2 and BIM were removed, and the (GGSG)_4_ linker was inserted between the C-terminus of BCL2 and the N-terminus of BIM_EL_ for structure prediction. (**D**) Binding curves for each BIM splice variant to BCL2 used for *K*_d_ fitting. All data were fitted with equation (1) in the Methods section. The binding curves accoring to prey proteins are individually displayed. Data are presented as means ± s.d. (*n* = 3). (**E**) Fitted *K*_d_ and 95% confidence intervals (CI) for PPIs between BCL2 and BIM splice variants. (**F**) Dynamic changes in *K*_d_ according to BIM splicing.

Next, to assess the specificity of the reconstituted target PPI reactions within the SMPC platform, we introduced mutations into conserved residues within the BH3 domain of BIM_EL_, the longest splice variant of BIM ^44-45^. Analysis revealed a progressive decline in PPI counts with increased evolutionary conservation of the residue, as evidenced by a marked decrease in interactions upon mutation of the highly conserved G156 residue (Figure 1G, H).^15, 17, 46-47^ This interaction was barely discernible from that of a negative control reaction (Figure 1H). This data underscores the pivotal role of the conserved BH3 domain in mediating the reconstituted PPIs between BCL2 and BH3-only proteins.

Significantly, the basic technical requirements for the SMPC assay—the overexpression of target proteins in mammalian cells—facilitates a versatile combination of baits and preys (Figure S4). This enabled the exploration of PPIs across a range of protein pairs extending beyond the BCL2 bait. We diversified our study by surface-immobilizing BCLxL- or MCL1-mCherry bait proteins as alternatives to BCL2, introducing a variety of BH3-only prey proteins. This strategic approach yielded PPI counts for an array of bait-prey configurations and revealed significant aspects of the BCL2 family’s PPI network, including the remarkably high-affinity interaction between BAD and BCLxL, which was conspicuously absent in the reactions involving MCL1 (Figure 1I).^12, 19^

### Profiling PPIs between BCL2 and splice variants of BIM

We utilized our SMPC platform to quantitatively ascertain the *K*_d_ values for multiple protein-protein interactions. To test the platform’s ability to distinguish between PPIs of closely related but distinct proteins, we evaluated several BIM splice variants (BIM_EL_, BIM_L_, BIM_S_, and BIM_XS_), which differ in their N-terminal exon domains preceding the BH3 domain (Figure 2A).^44-45^ According to the established structures of the BCL2-BIM complex, the BCL2 binding groove accommodates only the BH3 domain.^28^ AlphaFold2 predictions failed to provide a defined structure for these N-terminal domains of BIM alone or in complex with BCL2,^48-49^ hinting that areas other than the BH3 domain may be disordered even when BCL2 is present (Figure 2B, C, and Figure S5). Therefore, our comparison across the BIM variants aimed to understand the influence of these intrinsically disordered regions on PPI stability.

In optimizing the experimental setup to evaluate PPI strength between BCL2 and BIM variants, when using the BIM_EL_ variant with its transmembrane domain (TMD) intact, we noted that its PPI counts were subtly influenced by the type of detergents in the reaction buffer (Figure S6A). Consequently, we chose to analyze the PPIs of the TMD-truncated BIM variants in a buffer containing 0.03% Triton X-100, which not only increased the PPI count numbers per se and but also improved the consistency of our SMPC experiments across multiple repetitions (Figure S6B). We noted occasional cleavage of fluorescent tags from prey proteins (Figure S6C-E) but, aside from BIM_XS_, consistently found the remaining intact portion to exceed 60% for all variants assessed. This cleaved fraction was accounted for in our *K*_d_ calculations unless otherwise stated (and *K*_i_ values, see below).

When we immobilized a constant amount of BCL2-mCherry proteins (∼2,000 per field of view), we added the BH3 domain of BIM (BIM_BH3_) in increasing concentrations to determine PPI counts. These counts were converted into occupancy values, representing the ratio of BCL2-mCherry proteins bound by an eGFP-tagged prey protein. By plotting these occupancy values against the BIM_BH3_ concentrations, we deduced a *K*_d_ of 53.6±1.4 nM with an intra-assay coefficient of variance (CV) of approximately 6.6% (Figure 2D, E, and table S1).

In extending our analysis to additional BIM splice variants, we encountered an intriguing observation: the length of the splice variant did not correlate with a simple increase or decrease in occupancy value (Figure 2D-F). Instead, we observed complex patterns in the binding curves, indicative of the unique influence each exon domain has on BIM’s affinity for BCL2. The inclusion of the E2 domain (BIM_S_) significantly raised the occupancy values, thereby reducing the *K*_d_ from 53.6 nM to 15.7 nM (Figure 2D-F). Conversely, incorporating the E4 domain (BIM_L_ as opposed to BIM_S_) resulted in a marked decrease in occupancy, increasing the *K*_d_ to 63.3 nM (Figure 2D-F). These findings demonstrate that while the E2 domain promotes BCL2-BIM PPIs, the E4 domain undermines them. Introducing the E3 domain between E2 and E4 (BIM_EL_ versus BIM_L_) increased the binding curve, yielding a *K*_d_ of 27.3 nM, signifying that the presence of E3 considerably strengthens the BCL2-BIM interaction (Figure 2D-F).

Together, the SMPC platform enables swift PPI count determination for 40 different reactions, while demanding minimal experimental requirements. The resulting data sets reveal the individual contributions of domains to the BCL2-BIM interaction. Our data suggest that the BIM isoforms have varying and dynamic influences on apoptotic regulation through their differential affinities for BCL2 and other anti-apoptotic proteins.

### Profiling multi-body interactions between BCL2 family proteins and ABT-199

We subsequently leveraged our platform to investigate the impact of BCL2 mutations, G101V and D103E, identified in relapsed CLL patients post-ABT-199 therapy, on the complex interplay among BCL2, BH3-only proteins, and ABT-199. These mutations reside within the second alpha helix of BCL2, in a region critical for binding both the BH3 domain and ABT-199, suggesting they could modulate the affinity of BCL2 for its BH3-only protein ligands and for ABT-199 (Figure 3A).^35, 37^

**Figure 3.**
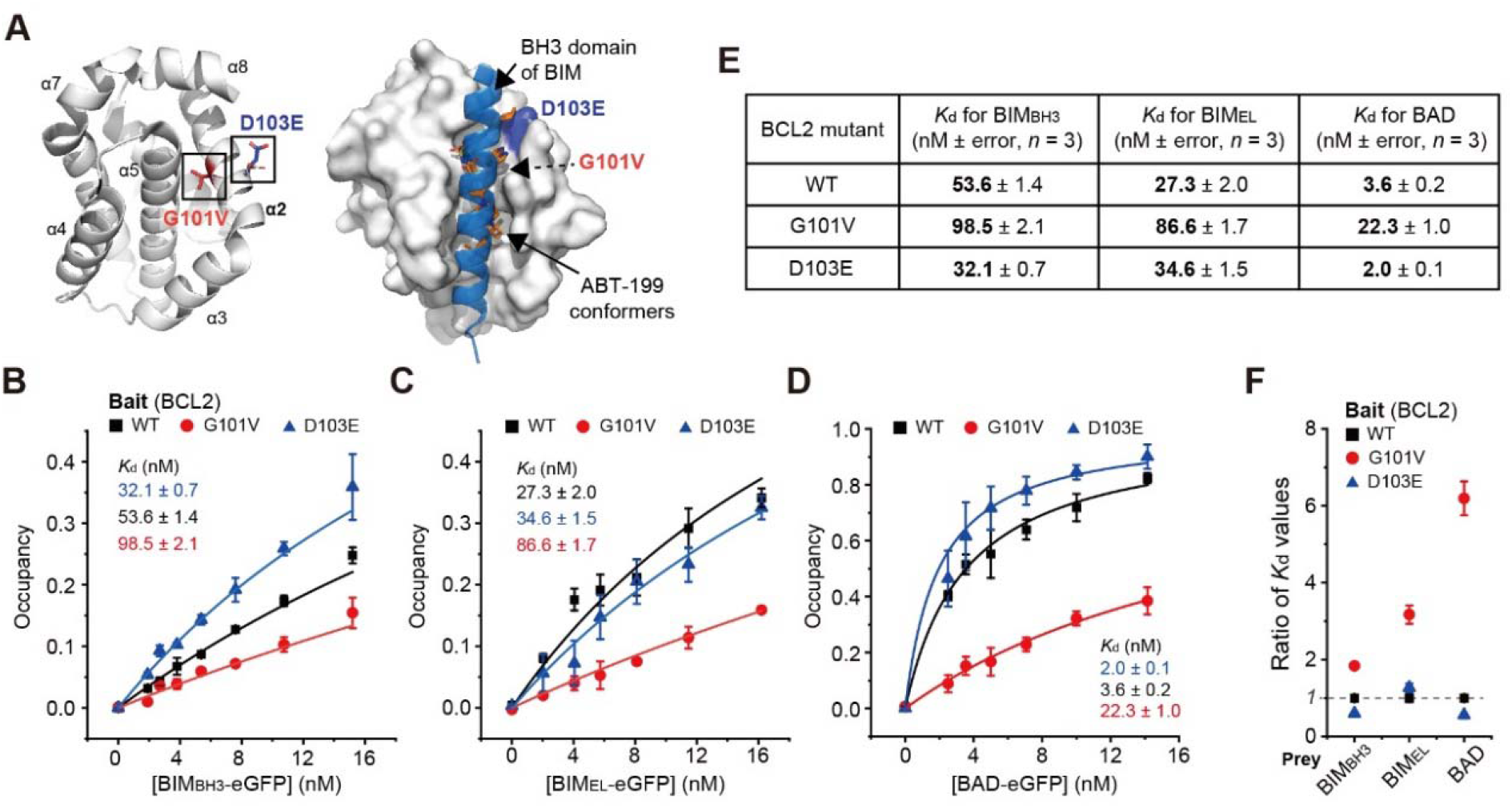
G101V and D103E mutants have opposite effects on BCL2’s interactions with BH3-only proteins. (**A**) Predicted BCL2 structures showing mutated residues (G101V and D103E) in comparison with the PDB structure (PDB id: 6O0K). BCL2 is shown in white, the BH3 domain of BIM is shown in blue, and ABT-199 conformers are shown in orange. G101 (V101) and D103 (E103) residues on the BCL2 are labeled. (**B**-**D**), Binding curves for each BH3-only protein to mutant BCL2s for *K*_d_ fitting. (B) BCL2-BIM_BH3_ PPIs, (C) BCL2-BIM_EL_ PPIs, and (D) BCL2-BAD PPIs. All data were fitted with equation (1) in the Methods section. Data are presented as means ± s.d. (*n* = 3). (**E**) Fitted *K*_d_ for PPIs between mutant BCL2s (WT, G101V, and D103E) and BH3-only proteins (BIM_BH3_, BIM_EL_, and BAD). The 95% CI values are described in Table S2. (**F**) Comparing the ratio of *K*_d_ values relative to BCL2_WT_ PPIs.

The reductionist’s nature of SMPC enabled us to reconstruct these intricate interactions incrementally, beginning with the BCL2 interactions with a suite of BH3-only proteins, followed by the introduction of the BH3 mimetic, ABT-199, to evaluate its impact on these interactions. As predicted by the steric properties of valine, and in agreement with existing literature, the G101V mutation notably disrupted the interactions between BCL2 and all tested BH3-containing proteins (BIM_BH3_, BIM_EL_, and BAD) (Figure 3B-F).^34-35^ The D103E mutation, however, conferred a modest yet significant increase in the binding affinity of BCL2 for BIM_BH3_, lowering the *K*_d_ from 53.6±1.4 nM (wild-type BCL2) to 32.1±0.7 nM (Figure 3B, E). These binding profiles were obtained by averaging the outcomes of three distinct experiments, thereby allowing us to discern relatively small shifts in *K*_d_ values (with an intra-assay coefficient of variation of 8.7%) (Tables S2 and S3). In assays with BIM_EL_, the D103E mutation resulted in a decreased affinity relative to wild-type BCL2. Hence, the presence of the D103E mutation unfavorably altered the interaction between BCL2 and BIM_EL_’s N-terminal disordered domains (E2 to E4), negating the enhanced binding observed with BIM_BH3_ and BCL2_D103E_ (Figure 2A, and Figure 3C, E). Employing full-length BAD as the prey molecule yielded analogous shifts in binding dynamics: G101V mutation resulted in substantial inhibition, whereas D103E mutation slightly but significantly strengthened the PPI (Figure 3D, E).

To quantify the effects of ABT-199 on the PPIs of each BCL2 mutant, we mixed ABT-199 with the BH3-only prey proteins under a set of designated conditions and counted the residual PPI complexes (Figure 4A). As anticipated, the more ABT-199 were included, the lower the residual PPI count, indicating competition among the prey molecules and ABT-199 for BCL2 binding. By fitting the inhibition curves of eight different ABT-199 concentrations, all with WT BCL2 as the bait, we were able to determine *K*_i_ values of 17.5 ± 1.0, 53.7 ± 4.8, and 4.4 ± 0.1 nM for BIM_BH3_, BIM_EL_, and BAD, respectively (Figure 4B-E). We further confirmed the specificity of these ABT-199 effects because the inhibition completely vanished when we switched the surface-immobilized bait to either BCLxL or MCL1 (Figure S7).

**Figure 4.**
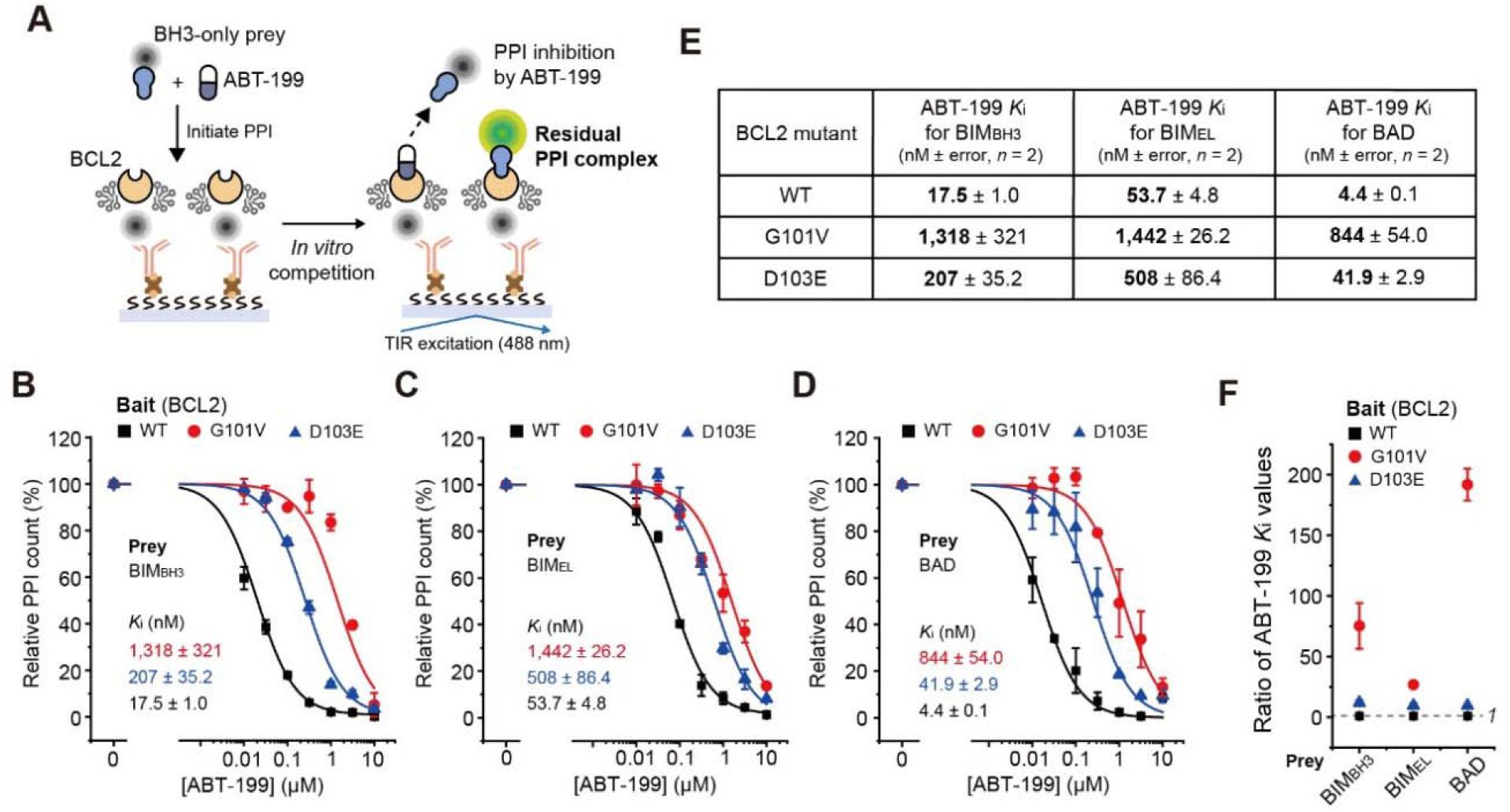
BCL2_G101V_ and BCL2_D103E_ take different routes to acquire resistance to ABT-199. (**A**) Schematic for the *in vitro* competition of ABT-199 with the PPIs between BCL2 and the BH3-only proteins. (**B**-**D**) Competitive ABT-199 inhibition curves for the residual PPI between each BH3-only protein to mutant BCL2s for *K*_i_ fitting. (B) BCL2-BIM_BH3_ PPIs, (C) BCL2-BIM_EL_ PPIs, and (D) BCL2-BAD PPIs. All data were fitted with equation (2) in the Methods section. (**E**) Fitted *K*_i_ for ABT-199 with each PPI pair. Each PPI signal is normalized to the corresponding ABT-199-untreated (0 μM) samples. Data are presented as means ± s.d. (*n* = 2). (**F**) Comparing the ratio of ABT-199 *K*_i_ values relative to BCL2_wt_ PPIs.

Together, our SMPC data provide a compelling account for why CLLs with this *BCL2* mutation acquired resistance to ABT-199 treatment. In the presence of the BCL2 mutations tested, we observed a consistent shift in the inhibitory curves towards higher ABT-199 concentrations, indicating a reduced affinity of the BCL2 mutants for ABT-199 (Figure 4B-E). Notably, the G101V mutation, which diminished the interactions of BCL2 with BH3-only protein partners, similarly attenuated the binding to ABT-199. This attenuation was quantified as elevations in the *K*_i_ values by multiples of 75, 25, and 190 for BIM_BH3_, BIM_EL_, and BAD, respectively (Figure 4F). Remarkably, these *K*_i_ value increments for the G101V mutation far exceeded those of the *K*_d_ values for the corresponding PPIs (compare Figure 3F with Figure 4F), suggesting that the mutation’s primary consequence is a diminished affinity for ABT-199 rather than an impaired ability to sequester BH3-only proteins. The D103E mutation exhibited a subtler decrease in ABT-199 affinity, with *K*_i_ values rising approximately tenfold for each of the three BH3-only proteins assessed (Figure 4B-F). While the D103E mutation maintained relatively high affinity for various pro-apoptotic proteins, it still showed significant increases in *K*_i_ values. An *in silico* exploration of the D103E variant was carried out to elucidate the structural basis for its differential impact on affinity for BIM_BH3_ and ABT-199 (Figure S8). Collectively, our SMPC data elucidate the mechanisms by which CLL cells harboring the BCL2 mutation develop resistance to ABT-199 therapy.

## Conclusion

Our investigation utilized the SMPC platform to elucidate the interactions between BCL2 and its mutants with an array of pro-apoptotic BH3-only proteins. The direct use of crude cell extracts is a distinguishing advantage of this platform, significantly simplifying the experimental setup in contrast to other methods for quantifying protein-protein interactions (PPIs). Our findings indicate that when target proteins are overexpressed, endogenous proteins minimally interfere with the target PPIs. This facilitates the omission of protein purification steps, which can be laborious for mammalian proteins, while maintaining post-translational modifications and avoiding the aggregation issues often encountered with membrane proteins and intrinsically disordered regions.

The integration of eGFP and mCherry into our methodology has strengthened its robustness and adaptability across diverse PPI studies. These fluorescent tags serve dual functions as reporters and IP handles. Due to the ultra-high affinity of some commercial antibodies for these tags, we achieved efficient IP within 15 minutes. Additionally, the absence of homologs for these fluorescent proteins in mammalian proteomes virtually eliminated nonspecific IP. A hurdle we encountered was the occasional proteolytic cleavage of the fluorescent tags from their conjugated proteins. Nevertheless, since the cleaved fraction typically represented less than 40% of the overall protein, we managed to adjust our calculated *K*_d_ and *K*_i_ values with less than a two-fold correction. Future efforts will focus on optimizing the linker sequence of these fluorescent tags to minimize such cleavage.

The imaging chips of our SMPC platform are designed for compatibility with standard 96- or 384-well plates, thus enabling parallel analysis of numerous PPIs. The reactions occur near equilibrium within individual reaction chambers, and the use of a single-molecule TIR fluorescence microscope equipped with auto-focusing enables PPI quantification by averaging the single-molecule eGFP signals over 300 ms. We typically compile over more than ten images per chamber to solidify our statistical confidence, rendering occupancy value determination for each condition completed within a minute—a significant time reduction compared to western blotting or surface plasmon resonance (SPR) analyses. Through this approach, we can derive *K*_d_ and *K*_i_ values for complex multi-component interactions.

The scope of our study encompassed 21 PPI pairs involving BCL2 and its mutants. This allowed for an in-depth analysis of the influence of the variably spliced N-terminal regions, which are primarily disordered, on the affinity of BIM for BCL2. Furthermore, we characterized the impact of BCL2 mutations found in relapsed CLL patients on the protein’s interaction with BH3-only proteins and the therapeutic agent ABT-199. Our research revealed that the mutations G101V and D103E affect BCL2’s interactions in notably distinct manners. While G101V impairs the sequestration of pro-apoptotic proteins by BCL2, it significantly disrupts BCL2’s affinity for ABT-199, elucidating the pronounced resistance to the drug observed in CLL cases harboring this mutation. Contrastingly, the D103E mutation either sustains or enhances the binding to BH3-only proteins when compared to the wild type but exhibits a marked decrease in ABT-199 binding. We anticipate that the SMPC platform will be instrumental in the future analysis of additional BCL2 mutations, particularly those in the second helix. In summary, the SMPC platform’s lower experimental demands, rapidity of measurement, and the quality and precision of its data output poise it to be a pivotal tool in the generation of extensive, quantitative PPI datasets.

## Supporting information

Supplementary Information includes Materials and Methods, Supplementary Figures 1-8, and Supplementary Tables 1-3.

## Acknowledgement

We thank Hongwon Lee, Byoungsan Choi, Kyung Chan Park, Hyunwoo Kim, and Shi Ho Kim for critical reading of the manuscript, and Changwon Kim for active discussion concerning data analysis. This work was supported by the National Grants for Leading Scientists (NRF-2021R1A3B1071354 to T.-Y.Y.) and the Bio Medical Technology Development Program (NRF-2018M3A9E2023523 to T.-Y.Y.) funded by the National Research Foundation of South Korea.

## Author Contribution

T.-Y.Y. conceived the project. C.C. and T.-Y.Y. designed the experiments. C.C. performed all experiments and analyzed data. C.C. performed data visualization. C.C. and T.-Y.Y. wrote the manuscript.

## Competing Interests

The authors declare no competing financial interest.

## Notes

### Competing Interest Statement

CC and T-YY filed a patent on these findings (10-2020-0157961).

