## Supplementary Information includes Materials and Methods, Supplementary Figures 1-8, and Supplementary Tables 1-3. for "Profiling multi-body interactions of BCL2 with single-molecule co-immunoprecipitation reveals the molecular mechanism of ABT-199 resistance"

**Antibodies**

Biotin anti-RFP antibody (ab34771; Abcam) was used to immobilize the mCherry-labeled proteins and to detect the aforelisted protein by western blot. Biotin anti-GFP antibody (ab6658; Abcam), MCL1 (94296S; Cell Signaling Technology), BCLxL (sc-56021; Santa Cruz Biotechnology), BCL2 (15071; Cell Signaling Technology), BIM (sc-374358; Santa Cruz Biotechnology), BAD (sc-8044; Santa Cruz Biotechnology), β-Tubulin (2128; Cell Signaling Technology), HRP-linked anti-rabbit IgG (7074S; Cell Signaling Technology), and HRP-linked anti-mouse IgG (7076S; Cell Signaling Technology) antibodies were used to detect the aforelisted proteins by western blot.

**Drug reagents**

ABT-199 (HY-15531; MedChemExpress) was used for *in vitro* competition. The concentration of the drug was diluted to 10 mM with DMSO and stored at -80 ℃.

**Cell line**

HEK293T cells were purchased from the ATCC.

**Cell culture and collection**

HEK293T cells were grown in DMEM (D6429; Sigma Aldrich) supplemented with 10% (v/v) FBS (26140-079; Gibco) and 100 μg/ml penicillin/streptomycin (15140-122; Gibco). HEK293T cells were cultured in a humidified incubator at 37 ℃, 5% CO2. All cultured cells were rinsed with cold DPBS (D8537; Sigma Aldrich) and were collected with a scraper (90020; SPL Life Sciences) in 1 ml of cold DPBS. The cell suspension was centrifuged at 500 g for 3 minutes at 4 ℃. The supernatants were discarded after centrifuge, and the cell pellets were stored at -80 ℃ by snap-freezing with liquid nitrogen.

**Constructions and expressions of mCherry/eGFP-labeled proteins**

All mCherry-labeled proteins (*BCL2* (HG10195-M; SinoBiological), *BCLxL* (HG10455-M; SinoBiological), and *MCL1* (HG10240-M; SinoBiological)) and eGFP-labeled proteins (*BIM_EL_* (HG13816-G; SinoBiological) and *BAD* (HG10020-M; SinoBiological)) were isolated from their respective human cDNA library and constructed by Gibson assembly. All cDNAs were cloned into *pCMV* vectors to generate and express BCL2 family proteins with mCherry or eGFP. The mCherry protein was cloned into the C-terminus of cDNAs of BCL2, BCLxL, and MCL1. For the BIM_EL_ and BAD, eGFP was cloned into the C-terminus of each cDNA. Mutations of BCL2 (G101V and D103E) were introduced by PCR using the corresponding primers. BIM splicing variants (BIM_S_ and BIM_XS_) and TMD-truncated BIM splicing variants (BIM_EL_, BIM_L_, BIM_S_, BIM_XS_, and BIM_BH3_) were isolated in respective domains from *BIM_EL_-eGFP* construct by PCR. For the purification of BAD-eGFP, the TEV cleavage sequence (EDLYFQS) and *HaloTag* sequence were cloned into the C-terminus of cloned *BAD-eGFP* by Gibson assembly.

The resulting plasmids were introduced into HEK293T cells through transient transfection by using linear polyethylenimine (PEI) (23966-100; Polysciences) following the manufacturer’s instructions. Typically, 10 μg of plasmid DNA was mixed with 35 μg of PEI in 1 ml of DMEM, and the mixture was introduced to 90 mm^2^ cell culture plate containing ~2$\times$10^6^ of HEK293T cells. Transfected cells were collected 18 hours after transfection and stored at -80 ℃.

**Cell lysis**

Collected cells were suspended with the lysis buffer (1% (v/v) detergent, 50 mM HEPES with pH 7.4, 150 mM NaCl, 10% (v/v) glycerol, 1 mM EDTA, 2% (v/v) protease inhibitor cocktail (P8340; Sigma Aldrich), 2% (v/v) phosphatase inhibitor cocktail 2 (P5726; Sigma Aldrich), and 2% (v/v) phosphatase inhibitor cocktail 3 (P0044; Sigma Aldrich)). Triton-X100 (X100; Sigma Aldrich), Glyco-diosgenin (GDN) (GDN101; Anatrace), Tween 20 (P2287; Sigma Aldrich), 3-([3-cholamidopropyl] dimethylammonio)-2-hydroxy-1-propanesulfonate (CHAPSO) (C4695; Sigma Aldrich), and n-dodecyl-β-D-maltoside (DDM) (D310; Anatrace) were used for lysis.

HEK293T cells transfected with eGFP-labeled prey proteins were lysed with Triton-X100 lysis buffer, and HEK293T cells transfected with mCherry-labeled bait proteins were lysed with GDN lysis buffer. After lysis, the cell suspension was centrifuged at 15,000 g for 10 minutes at 4 ℃. The supernatants were isolated after centrifuge. The total protein concentration in each supernatant was measured with a DC protein assay kit (5000113, 5000114, 5000115; Bio-Rad) following the manufacturer’s instructions. The total concentration of the fluorescence proteins in each supernatant was measured with a Sense microplate reader (425-301; HIDEX). To quantify the fluorescence proteins, the laser wavelength of 488 nm and 544 nm were used for eGFP and mCherry excitation respectively. The calibration curves were prepared using the emission patterns from purified eGFP or mCherry proteins. The supernatants were then aliquoted and stored at -80 ℃ after snap-freezing with liquid nitrogen.

**Protein purification**

The HEK293T cells transfected with *BAD-eGFP-HaloTag* plasmids were suspended in purification lysis buffer (0.2% (v/v) Triton-X100, 50 mM HEPES with pH 7.4, 150 mM NaCl, 10% (v/v) glycerol, 1 mM EDTA, 2 mM TCEP (C4706; Sigma Aldrich), 1% (v/v) protease inhibitor cocktail, 1% (v/v) phosphatase inhibitor cocktail 2, and 1% (v/v) phosphatase inhibitor cocktail 3). The suspended cells were lysed by sonification and centrifuged at 15,000 g for 10 minutes at 4 ℃. The BAD-eGFP proteins in supernatant were purified with the HaloTag mammalian protein purification system (G6790; Promega) following the manufacturer’s instructions. The supernatant was nutated with HaloLink resin for 18 hours at 4 ℃. The resin was washed with purification lysis buffer and incubated with TEV elution buffer (1:50 HaloTEV protease, 50 mM HEPES with pH 7.4, 150 mM NaCl, 10% (v/v) glycerol, 1 mM EDTA) for 18 hours at 4 ℃. The eluate was loaded onto an Amicon ultra-15 centrifugal filter (UFC9030; Millipore) pre-equilibrated with purification lysis buffer. Only the BAD-eGFP fraction was concentrated on the filter, and the concentrate was then aliquoted and stored at -80 ℃ after snap-freezing with liquid nitrogen.

The purified proteins were confirmed by 6-15% gradient SDS-page gels (Figure S3). The concentration of the purified BAD-eGFP was quantified from the eGFP concentration measured by the Sense microplate reader.

**Western blot**

All cell extracts used for western blot were produced by the steps described in the cell lysis subsection. The loading concentration of each sample was quantified based on either total protein concentration or total mCherry/eGFP concentration. All samples were heated at 95 ℃ for 15 minutes with SDS containing sample buffer (EBA-1052; EPLIS Biotech), resolved with 6-15% gradient SDS-page gels, and transferred to PVDF membranes (IB401001; Thermo Fisher Scientific). PVDF membranes were blocked in 5% (w/v) skim milk for 1 hour at room temperature, and each membrane was immunoblotted with an appropriate primary antibody with 1% (w/v) BSA. After overnight incubation at 4 ℃, membranes were immunoblotted with an appropriate secondary antibody for 1 hour at room temperature. Protein bands were detected using an ECL (W3653-020; GenDEPOT) imaging system of ImageQuant LAS 4000 mini, and the band intensity was analyzed with ImageJ 1.53a.

**Preparation of PEG-coated coverslip for SMPC**

PEG-coated coverslip was produced using the previously reported methods.^1^ Briefly, the coverslip (48393-251; VWR) was rinsed with acetone for 15 minutes. After that, the coverslip was washed with 1 M KOH solution for 15 minutes. The coverslip was cleaned for 30 minutes with piranha solution (H_2_SO_4_:30% (w/v) H_2_O_2_ = 2:1). Amino silane (104884; Sigma Aldrich) was then immobilized onto the hydroxyl group of the cleaned coverslip surface by silanization reaction with silane solution (3% (v/v) Amino silane, 5% (v/v) acetic acid in methanol). After that, biotin-PEG (BIO-PEG-SVA-5K; LaySan Bio) and mPEG (mPEG-SVA-5K; LaySan Bio) powder were dissolved in buffer (0.1 M NaHCO_3_, 0.4 M K_2_SO_4_) to make PEG solution (Biotin-PEG:mPEG = 3:100, 100 mg/ml of PEG mixture in solution). The PEG solution was filtered using a 0.2 μm pore filter. PEG-coating was performed by spraying 80 μl of the filtered PEG solution on one side of the coverslip, covering the other coverslip, and incubated for 3 hours in a dark room at room temperature. The PEG-coated coverslip stack was separated and washed with distilled water. After that, the coverslips were dried with pure nitrogen gas. All coverslips were stored at -20℃ in a vacuum condition. Before the experiment, PEG-coated coverslips were equilibrated to room temperature for 10 minutes to avoid water condensation.

**Construction of SMPC imaging chip**

The dimension of the acrylic frame and the double-sided tape were customized to construct the imaging chip. The acrylic plates were drilled with 40 holes (3 mm diameter each) with 1 mm spacing in an array of 8 by 5. The drilled acrylic plates were molded into individual acrylic frames (60 mm×24 mm×3 mm). Each hole serves as an individual reaction chamber. The dimension of the double-sided tape was also customized to fit onto the acrylic frame.

To assemble the imaging chip, the double-sided tape was gently applied on top of the acrylic frame. Then, the PEG-coated coverslip was attached to the sticky side of the acrylic frame to form an imaging chip with 40 reaction chambers.

**SMPC**

Surface-immobilization: 5 μg/ml of Neutravidin (31000; Thermo Fisher Scientific) was loaded into each reaction chamber of the imaging chip and incubated for 10 minutes. The imaging chip was washed with Tx100-buffer (0.1% (v/v) TritonX-100, 50 mM HEPES with pH 7.4, 150 mM NaCl) to remove any unbound Neutravidin. Biotin Anti-RFP antibodies were loaded to each reaction chamber with 1:200 in Tx100-buffer and incubated for 10 minutes. The imaging chip was washed with GDN-buffer (0.01% (w/v) GDN, 50 mM HEPES with pH 7.4, 150 mM NaCl, 1% (v/v) Glycerol, 1 mM EDTA). After washing, crude cell extracts were diluted based on the concentration of mCherry labeled bait proteins in extracts and were incubated for 15 minutes. The imaging chip was washed with the GDN-buffer, and was mounted on the TIR fluorescence microscope to measure the fluorescence signals from surface-immobilized mCherry without the removal of GDN-buffer.

**PPI reaction:** After lysis of the HEK293T cells expressing eGFP-labeled prey proteins, the crude cell extracts were diluted to appropriate eGFP concentration (10 nM), *C*_tot_ (less than 0.1 mg/ml typically), and 0.03% (v/v) of Triton X-100 concentration in the final extracts. The final extracts were incubated for 10 minutes after surface immobilization of bait proteins. The imaging chip was placed on the TIR fluorescence microscope to measure the amount of PPI complexes without the removal of final extracts.

Based on the measured data, *K*_d_ of each PPI pair was fitted using the following equation (1) in OriginPro 2022.

$$V=V_{0}+\frac{V_{max}*[prey]}{K_{d}+[prey]} \left( 1 \right)$$

$V$=measured occupancy (count of PPI complexes/count of immobilized bait proteins), $V_{0}$=fixed initial occupancy (typically $V_{0}=0$), $V_{max}$=maximal occupancy (fixed at $V_{max}=1$), $\left[ prey \right]$=concentration of prey proteins, $K_{d}$=fitted dissociation constant.

***In vitro* competition:** In the process of dilution of extracts containing eGFP-labeled prey proteins, ABT-199 was mixed with the final extracts. The final extracts were incubated for 10 minutes after surface immobilization of bait proteins. The imaging chip was placed on the TIR fluorescence microscope to measure the amount of PPI complexes without the removal of final extracts.

Based on the measured data, the *K*_i_ of ABT-199 for each PPI pair was fitted using the following equation (2) in OriginPro 2022.

$$V=V_{0}+\frac{V_{max}-V_{0}}{1+\frac{K_{d}*(K_{i}+\left[ ABT \right])}{K_{i}*\left[ prey \right]}} (2)$$

$V$=measured normalized PPI, $V_{0}$=fixed minimum PPI (typically $V_{0}=0$), $V_{max}$= maximal PPI, $\left[ ABT \right]$=concentration of ABT-199, $\left[ prey \right]$= concentration of prey proteins, $K_{d}$=fixed dissociation constant measured in equation (1), $K_{i}$=fitted inhibitory constants of ABT-199.

**Detection of fluorescence by TIR microscope and post-image analysis:** All single-molecule fluorescence signals were measured using the home-built objective-based TIR fluorescence microscope as pre-reported methods.^1^ For TIR illumination, the laser wavelength of 488 nm and 532 nm were used for eGFP and mCherry excitation respectively. The single-molecule fluorescence signal was detected via the electron-multiplying charge-coupled device (EMCCD) (iXon 897, DU-897U-CS0-EXF; ANDOR) for 10 frames (typically 100 ms exposure) and recorded in a TIFF stack file format. Within each chamber of the imaging chip, ~12 images were taken from different locations in a spiral motion by the piezo controller (MS-2000-500-CP; ASI). The final images were carefully selected to avoid any biased analysis by filtering any photobleached and aggregated large dye cluster images. The number of single-molecule fluorescence counts and summation of total intensity in a TIFF stack were measured by MATLAB 2020a. A detailed protocol for counting single-molecule fluorescence signals is described in reference.^2^

**Structural analyses**

All structures (BCL2, BCL2_G101V_, BCL2_D103E_, BIM_EL_, BCL2-BIM_BH3_ complex, and BCL2-BIM_EL_ complex) were predicted by Alphafold2 software provided by ColabFold.^3^ The predicted lDDT scores of predicted structures were also obtained from ColabFold. For the BCL2-BIM_BH3_ complex was compared to the PDB structures (PDB id: 6O0k)^4^ for *in silico* analysis. Visualization of structural information was performed using PyMolwin software.

**Supporting Figures**

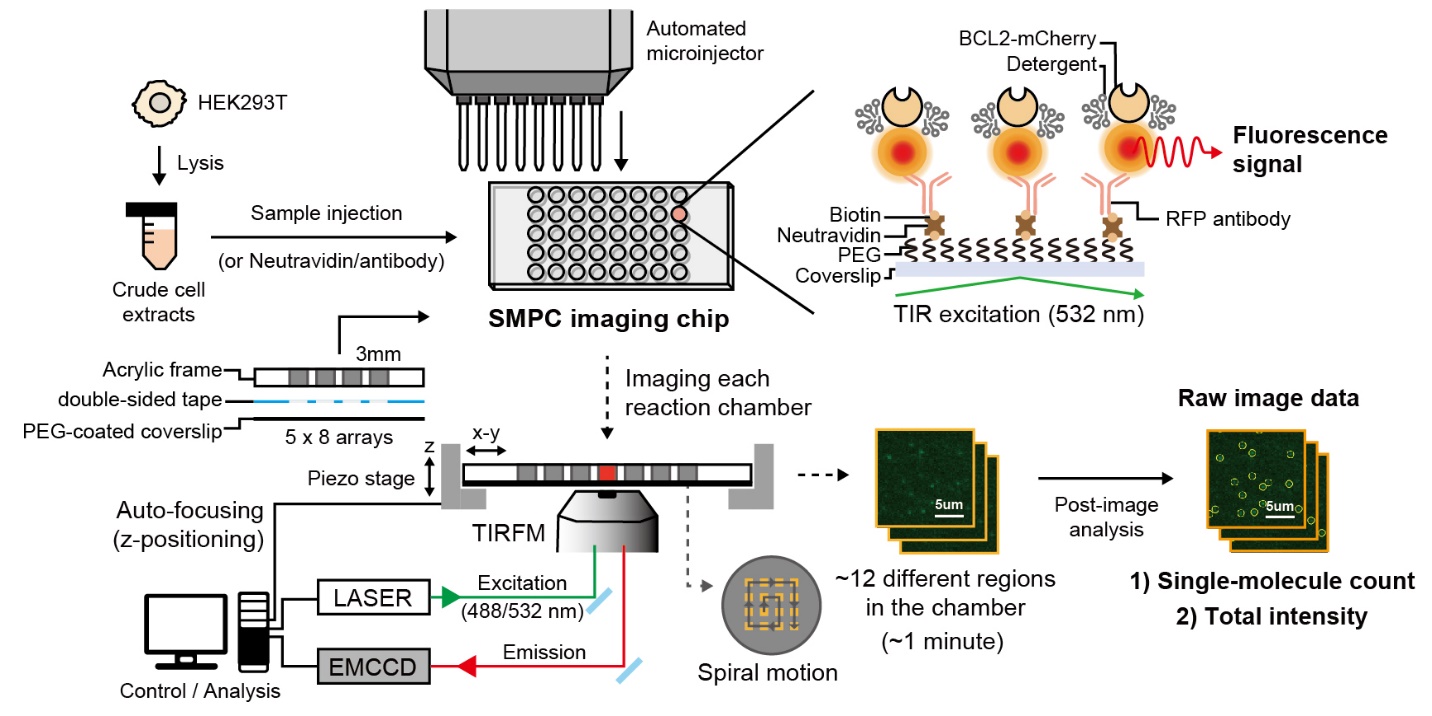

**Figure S1. Overview of instrumental setup and analysis strategy of SMPC.** Schematic of the SMPC with TIR fluorescence microscope (TIRFM) setup and analysis for counting fluorescence signals of immobilized proteins and PPI complexes. The SMPC imaging chip was assembled using double-sided tape with an acrylic frame and a PEG-coated coverslip. An automated microinjector was used to inject Neutravidin, antibody, and crude cell extracts into the reaction chamber. After surface-immobilization or PPI reaction, approximately 12 independent images were taken. Target single-molecule fluorescence counts (single-molecule count) and the total intensity data within the field of view (50×100 μm^2^) of the raw image data were obtained through computational post-image analysis. The detailed protocol is described in the Method section.

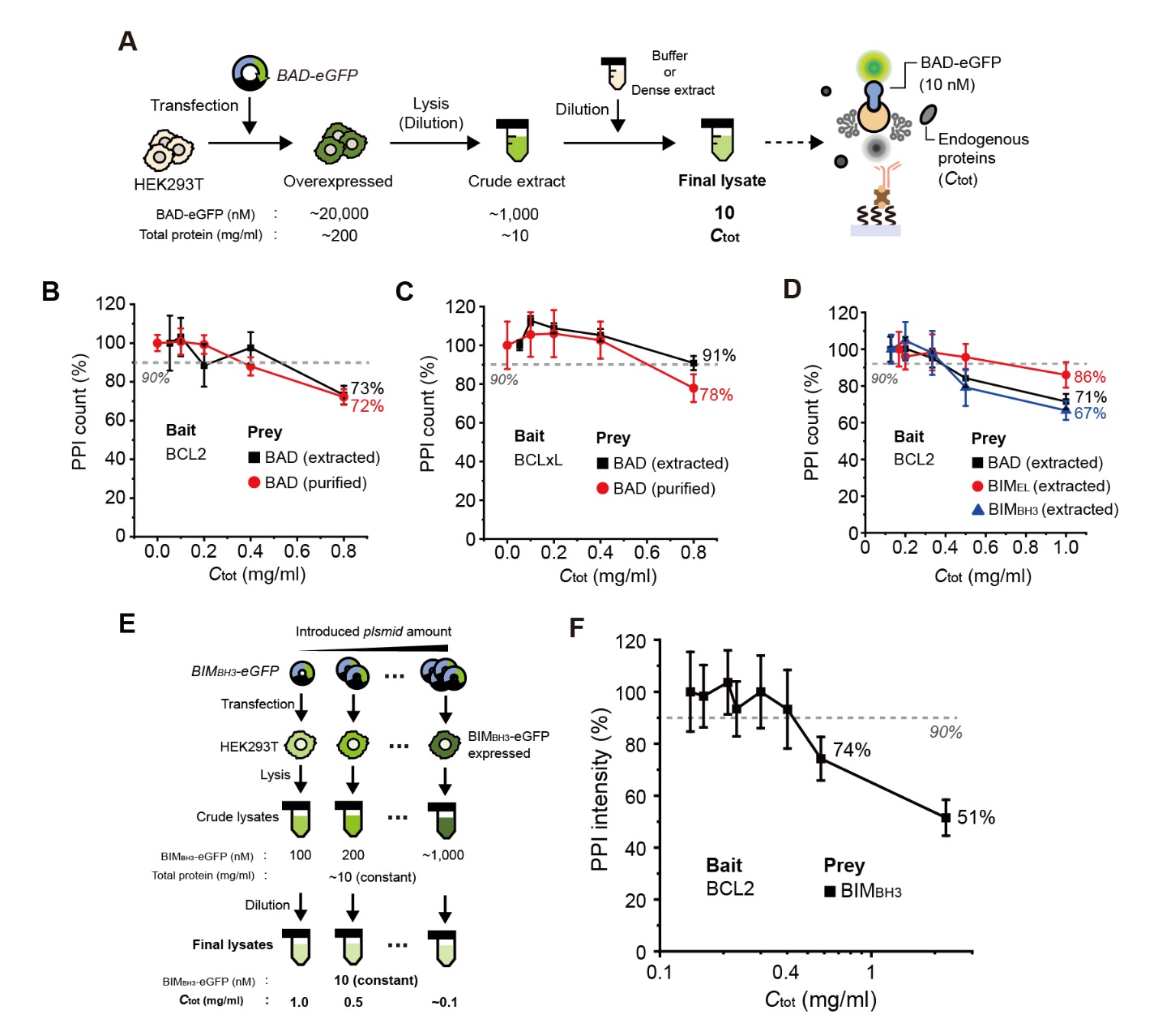

**Figure S2. Overexpression of PPI prey proteins for use in SMPC.** (A) Schematic of the process of diluting extracts from HEK293T cells overexpressing eGFP labeled BAD proteins to acquire the working concentration of 10 nM. (B-D) Relative changes in PPI counts with increasing *C*_tot_. (B) PPIs between BCL2 and BAD proteins. (C) PPIs between BCLxL and BAD proteins. (D) PPIs between BCL2 and BH3-only proteins in crude extracts (BAD, BIM_EL_, and BIM_BH3_). (E) Schematic for the expression of BIM_BH3_-eGFP with different *C*_tot_ values at the cellular level. (F) Relative change in BCL2-BIM_BH3_ PPI intensity according to the increase of *C*_tot_. All data are normalized to the lowest *C*_tot_ sample in each experiment.

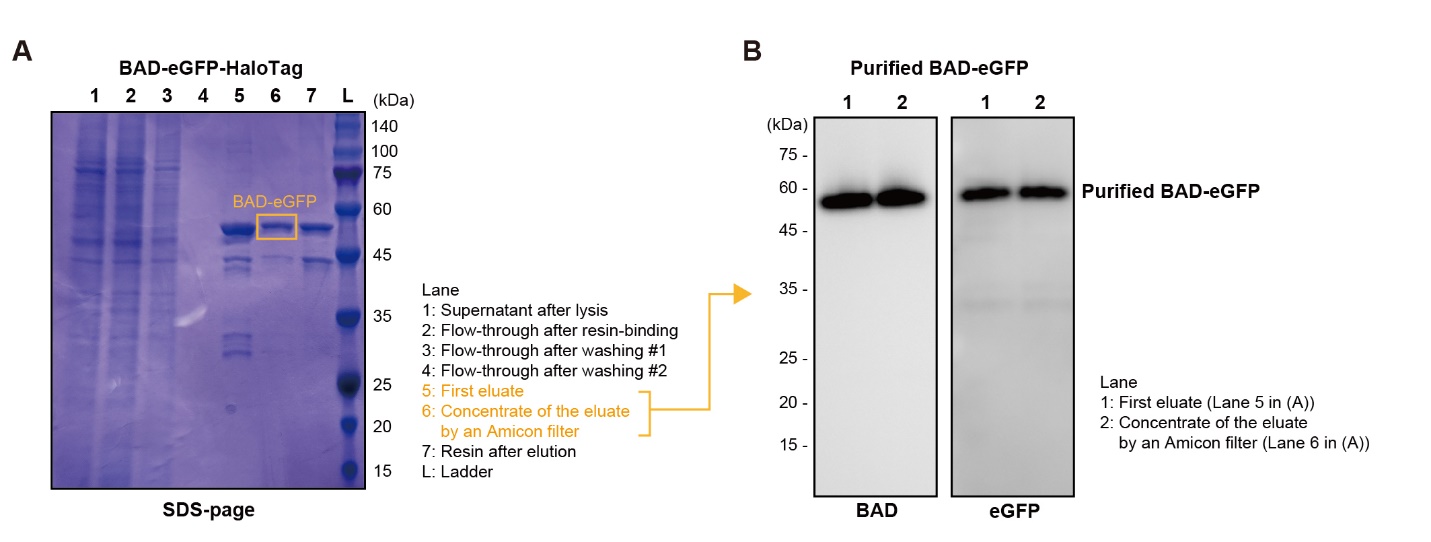

**Figure S3. Purification of the BAD-eGFP proteins from crude HEK293T extracts.** (A) SDS-PAGE analysis of the BAD-eGFP proteins purified from crude HEK293T extracts. (B) Western blot analysis of the purified BAD-eGFP proteins. The first eluate (Lane 5 in panel (A)) and the concentrate of the eluate by an Amicon filter (Lane 6 in panel (A)) were loaded respectively. The purified BAD-eGFP proteins were detected by anti-BAD antibody and biotinylated anti-GFP antibody.

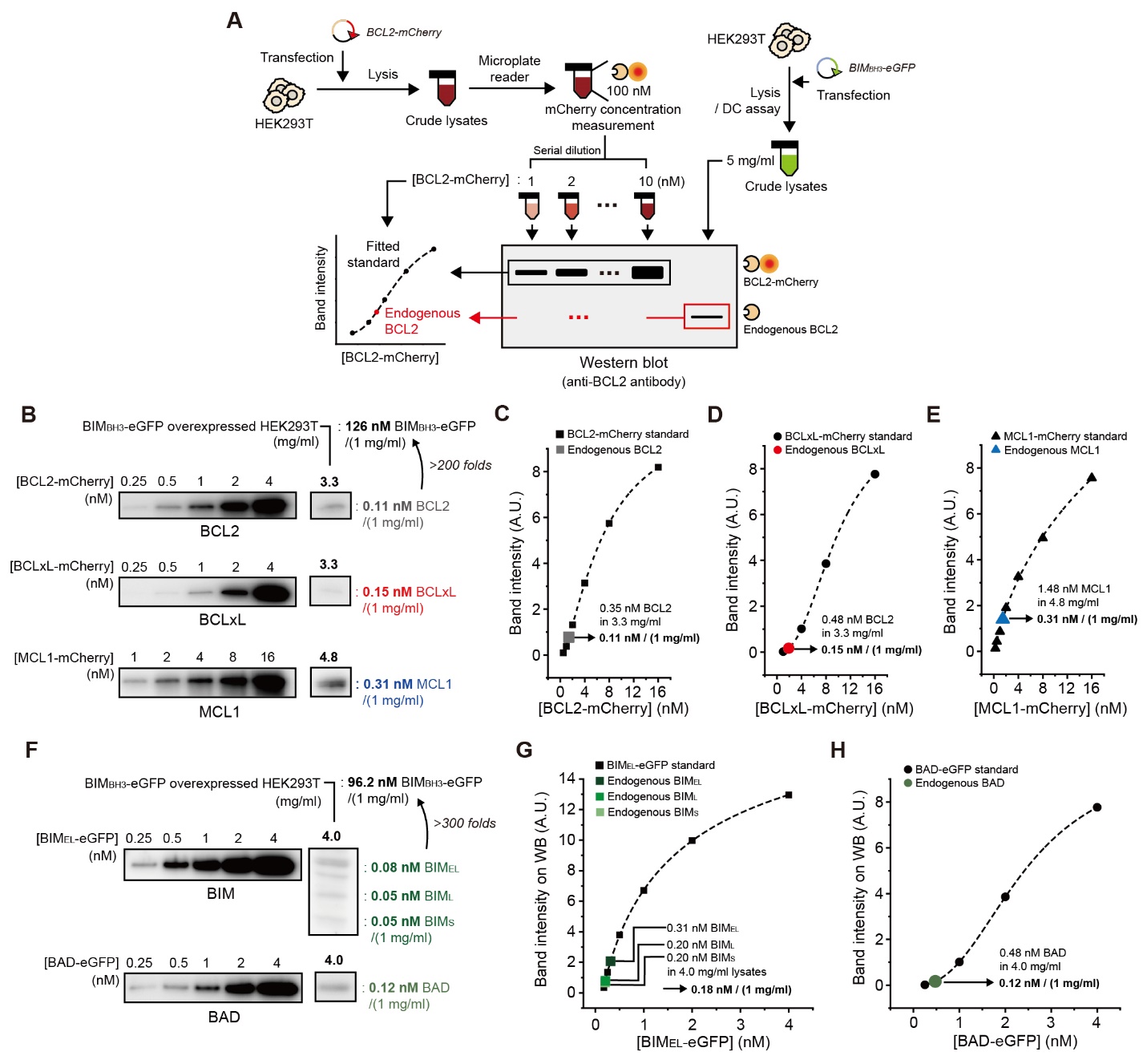

**Figure S4. Measurement amount of endogenous BCL2 family proteins in HEK293T extracts.** (A) Schematic for the measurement of endogenous BCL2 family proteins using western blot. mCherry-labeled BCL2 was overexpressed in HEK293T, and BCL2-mCherry concentration in crude cell extracts was quantified with a microplate reader. The extracts were serially diluted according to the mCherry concentration and loaded to the blot. The blot was detected with anti-BCL2 antibody, and the standard curve was fitted using the band intensity and the concentration of BCL2-mCherry. On the same blot, BIM_BH3_-eGFP overexpressed HEK293T cell extracts were loaded, and the level of endogenous BCL2 was estimated from the standard curve. (B) The concentration of endogenous anti-apoptotic BCL2 family proteins (BCL2, BCLxL, and MCL1) in HEK293T cell extracts after BIM_BH3_-eGFP overexpression. (C-E), Fitted standard curves to estimate the concentration of endogenous anti-apoptotic proteins in (B). (C) BCL2, (D) BCLxL, and (E) MCL1. (F) The concentration of endogenous BH3-only proteins (BIM and BAD) in HEK293T cell extracts after BIM_BH3_-eGFP overexpression. (G and H) Fitted standard curves to estimate the concentration of endogenous BH3-only proteins in (F). (G) BIM variants (BIM_EL_, BIM_L_, and BIM_S_), (H) BAD.

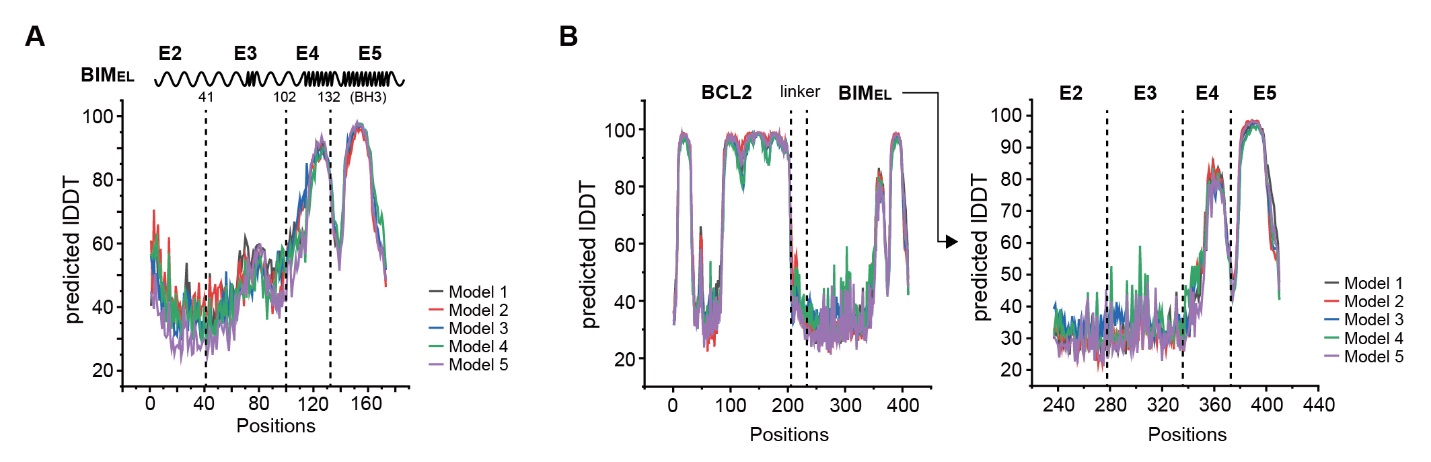
**Figure S5. Prediction of BIM_EL_ and BCL2-BIM_EL_ complex structure by AlphaFold2.** (A) Predicted the local Distance Difference Test (lDDT) scores of top five BIM_EL_ structure models. The spring-like figure depicts the part with the α-helix structure, and the rest for intrinsically disordered regions. The highest scored model is displayed in Figure 2B. (B) Predicted lDDT scores of top five BCL2-BIM_EL_ complex structure models. The right graph displays scores of BIM_EL_ positions only. The highest scored model is displayed in Figure 2C. TMD sequences of BCL2 and BIM were removed, and the (GGSG)_4_ linker was inserted between the C-terminus of BCL2 and the N-terminus of BIM_EL_ for complex structure prediction (A and B).

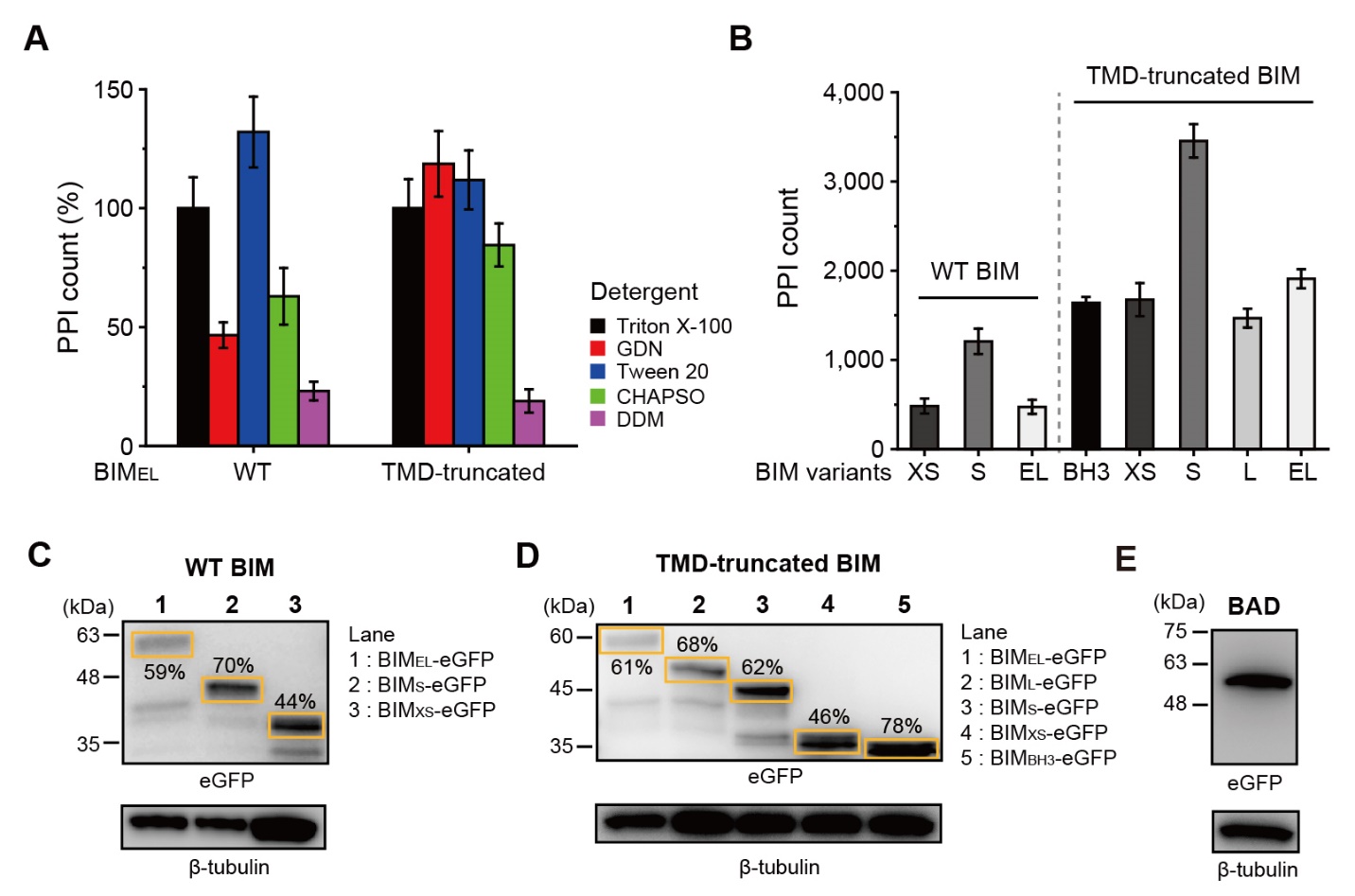

**Figure S6. The transmembrane domain of BIM was delicately affected to the detergent condition for PPI.** (A) Normalized relative changes in PPI counts of WT BIM_EL_ and TMD-truncated BIM_EL_ for BCL2 with various detergent conditions in the reaction buffer. The concentration of detergent in the reaction buffer was all maintained at 0.03%. All data are normalized to Triton X-100 conditions in each experiment. (B) PPI counts of 10 nM BIM variants for BCL2 in Triton X-100 condition. (C and D) Expression of splicing variants of BIM proteins in HEK293T. (C) WT BIM (BIM_EL_, BIM_S_, and BIM_XS_), (D) TMD-truncated BIM (BIM_EL_, BIM_L_, BIM_S_, BIM_XS_, and BIM_BH3_). (E) Expression of BAD-eGFP in HEK293T. 10 μg of crude cell lysates were loaded, and BH3-only proteins were detected by biotinylated anti-GFP antibody. (C-E) The band of the target protein and sporadic cleavage rate were indicated on blot images.

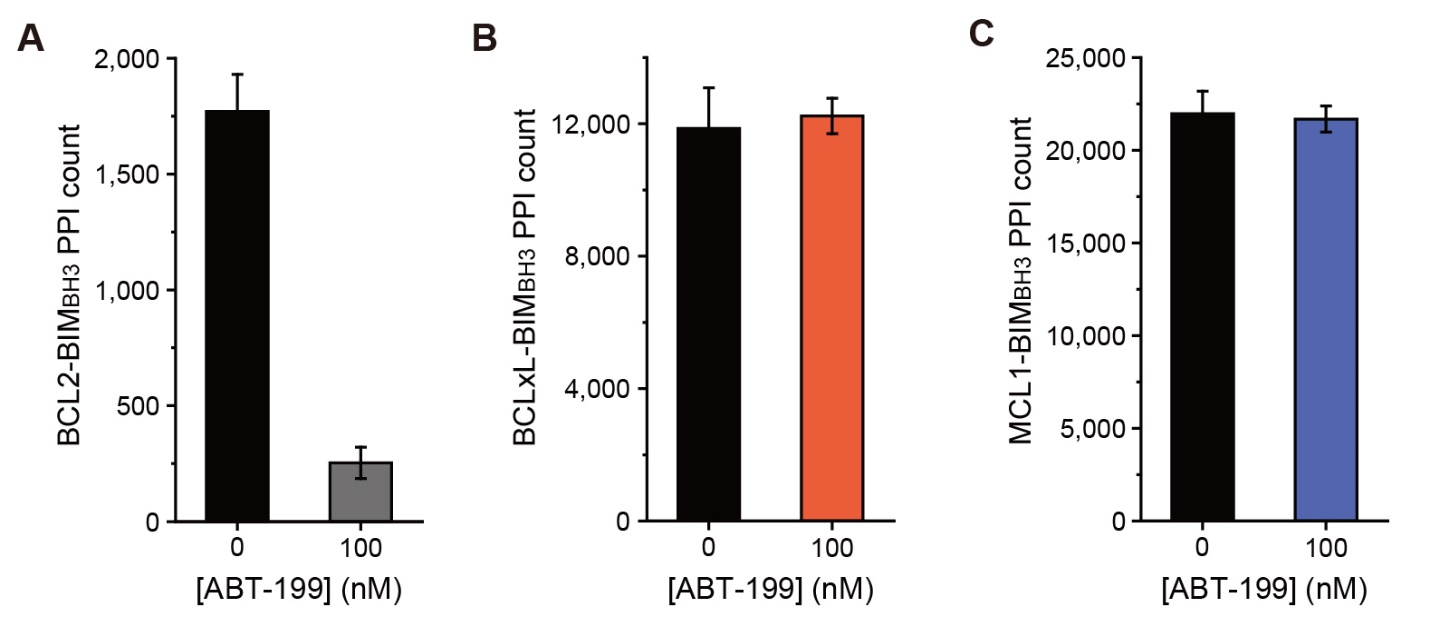

**Figure S7. ABT-199 selectively inhibited BCL2-BIM_BH3_ PPI.** (A-C), *In vitro* competition of ABT-199 for PPI between different surface baits (BCL2, BCLxL, or MCL1) and 10 nM of BIM_BH3_-eGFP preys. ABT-199 was mixed with BIM_BH3_-eGFP lysates before PPI reaction. (A) BCL2-BIM_BH3_ PPI, (B) BCLxL-BIM_BH3_ PPI, and (C) MCL1-BIM_BH3_ PPI.

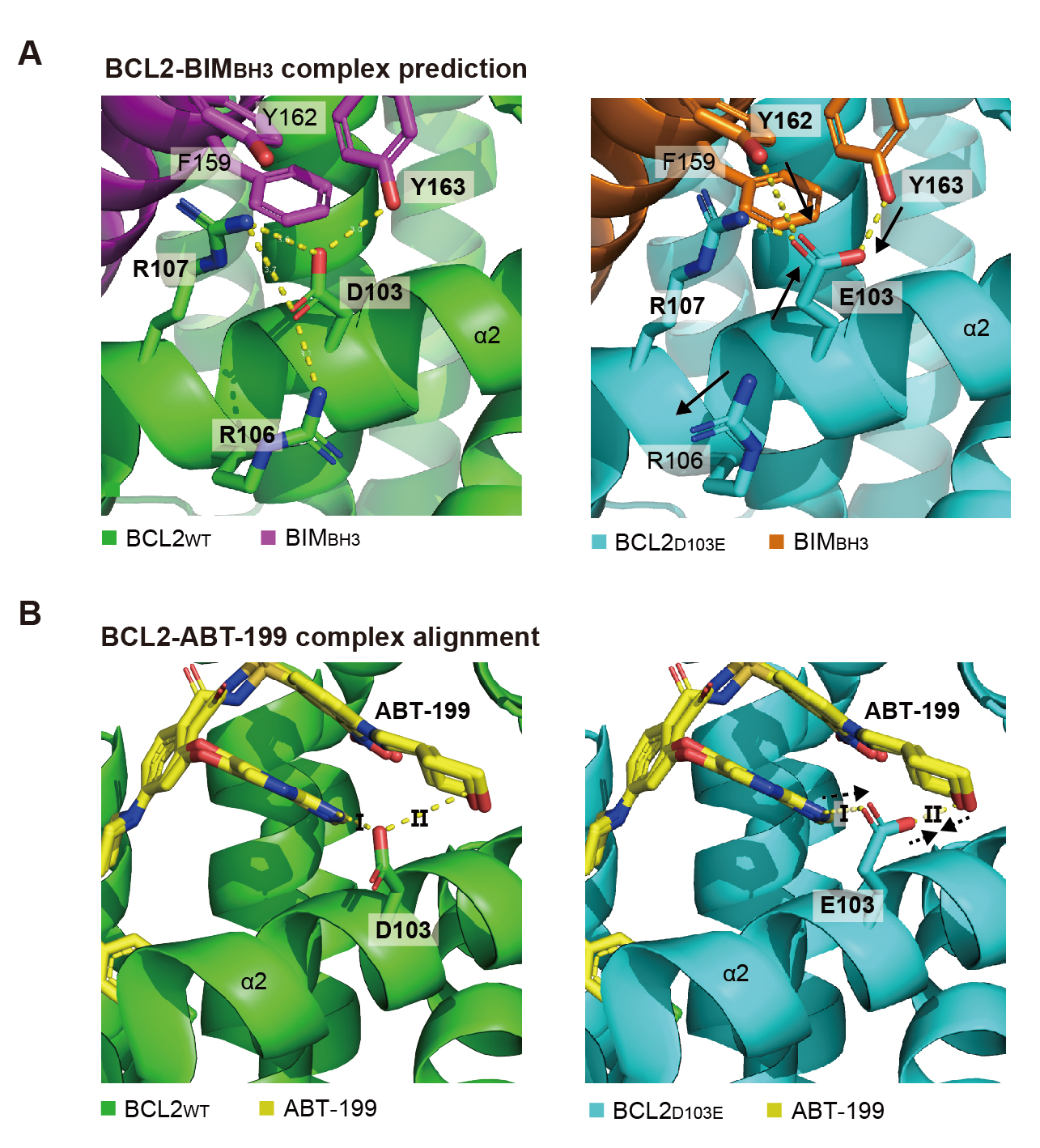

**Figure S8. *In silico* analysis of BCL2_D103E_ complex structures with BIM and ABT-199 by AlphaFold2**. (A) Comparison of the predicted structure of BCL2_WT_-BIM_BH3_ and BCL2_D103E_-BIM_BH3_ complexes. In E103, the negatively charged side chain is truncated in the opposite direction of WT (D103). The yellow dotted line indicates the interaction within 4Å, and the black arrow indicates the direction in which the interaction with each side chain changes after mutation (D103E). (B) Comparison of the predicted structure of BCL2_WT_-ABT-199 and BCL2_D103E_-ABT-199 complexes. The BCL2-BIM_BH3_ complex predicted in (A) and the known BCL2_G101V_-ABT-199 structure (PDB id: 6O0k) were aligned for comparison. The ABT-199 conformers are indicated in yellow. Interaction I refers to the electrostatic interaction between the indol-ring of ABT-199 and the side chain of D103 residue on BCL2. Interaction II refers to the hindrance between ABT-199 and D103 residue. E103 shows weaker electrostatic interaction for interaction I, and stronger hindrance for interaction II.

**Supporting Tables**

**Table S1. Fitted *K*_d_ of BIM variants for BCL2 to evaluate intra-assay CV.** All the *K*_d_ data are shown for each independent experiment. CV for each PPI pair was calculated by CV= s.d./mean (%) for three technically independent replicates. The intra-assay CV is the average of five CV values from each PPI pair.

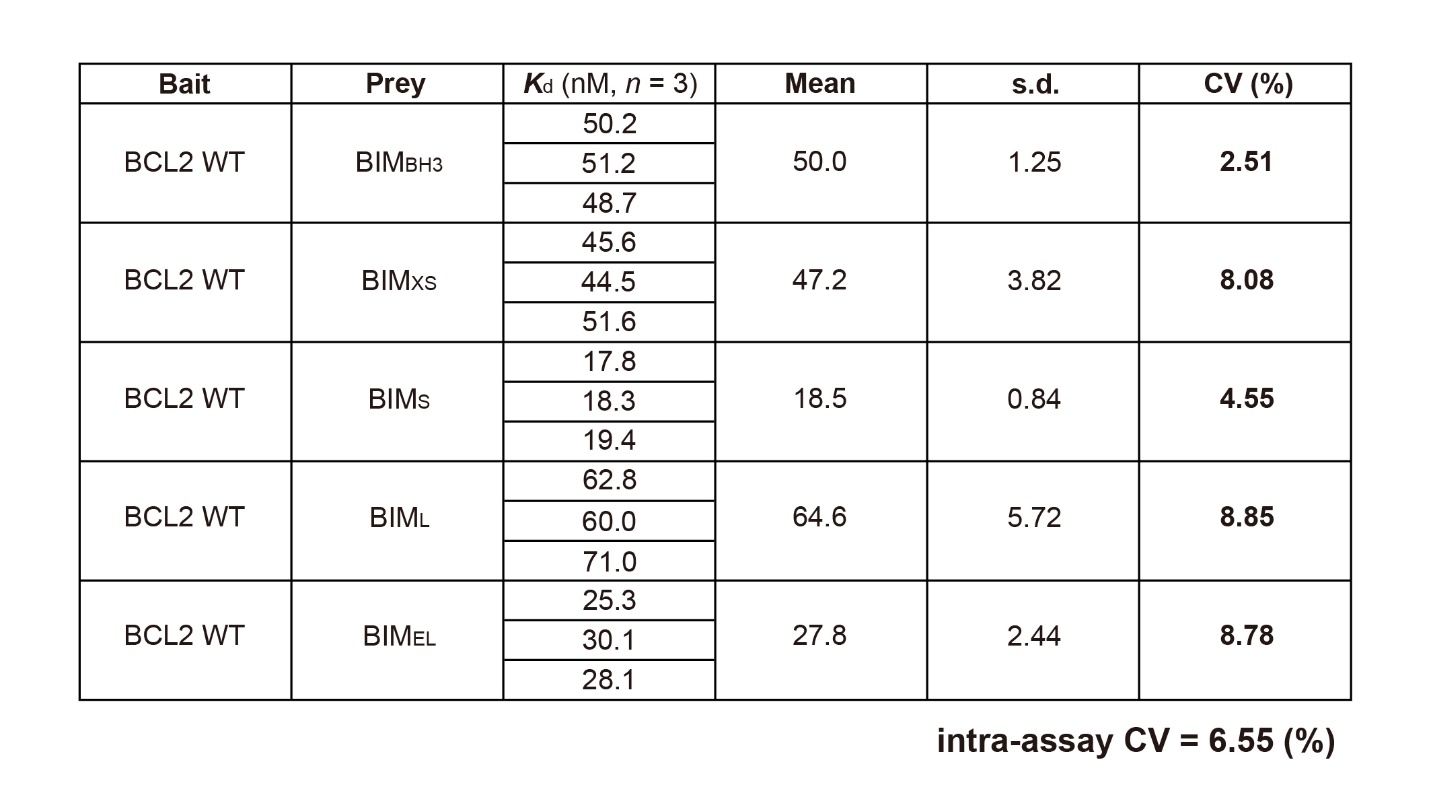

**Table S2. Fitted *K*_d_ and 95% confidence interval (CI) of BH3-only proteins for BCL2 mutants.** The fitted *K*_d_ of BH3-only proteins (BIM_BH3_, BIM_EL_, and BAD) for BCL2 mutants (WT, G101V, and D103E). All the *K*_d_ data shown were fitted by averages of three technically independent replicates.

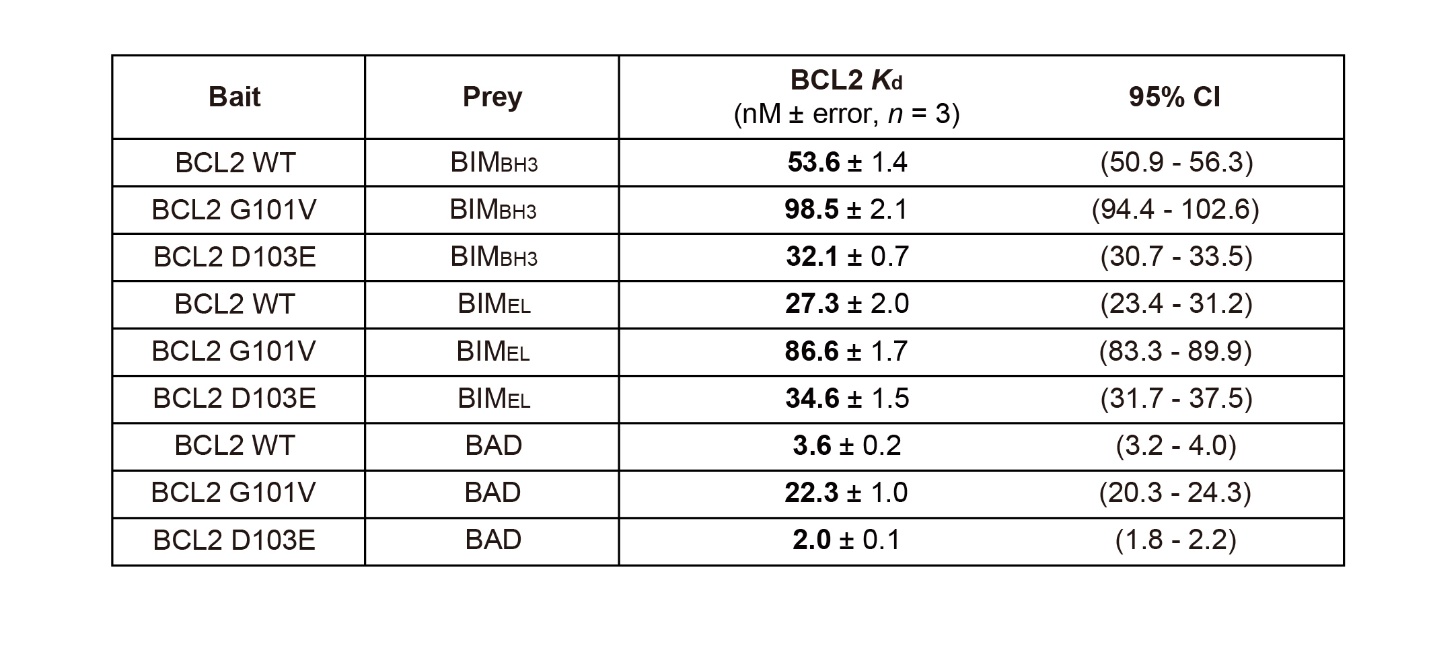

**Table S3. Fitted *K*_d_ of BH3-only proteins for BCL2 mutants to evaluate intra-assay CV.** The fitted *K*_d_ of BH3-only proteins (BIM_BH3_, BIM_EL_, and BAD) for BCL2 mutants (WT, G101V, and D103E). All the *K*_d_ data are shown for each independent experiment. CV for each PPI pair was calculated by CV= s.d./mean (%) for three technically independent replicates. The intra-assay CV is the average of five CV values from each PPI pair.

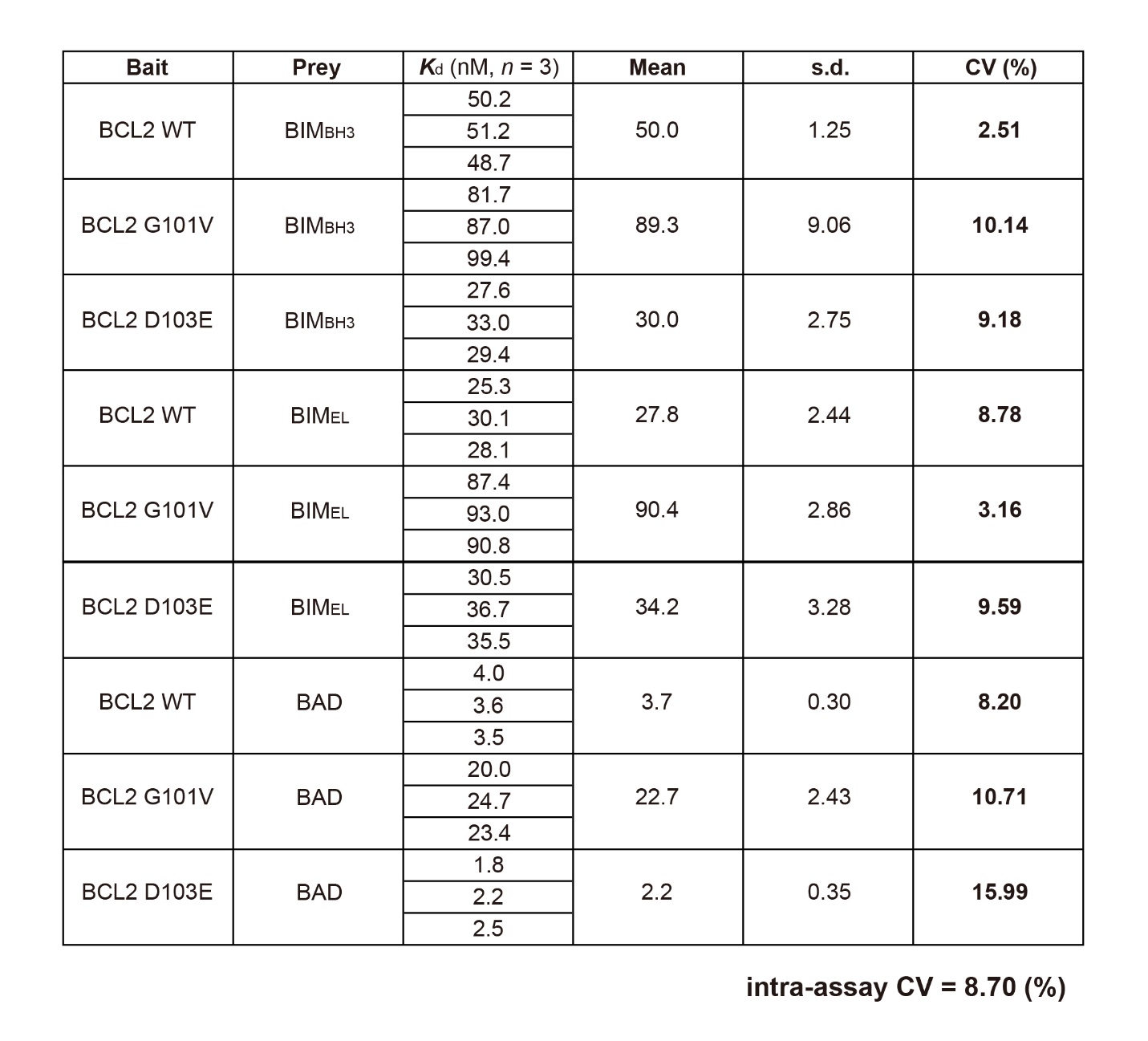
